# Sensitivity and Frequency Response of Biochemical Cascades

**DOI:** 10.1101/2023.09.14.557821

**Authors:** Michael A. Kochen, Joseph L. Hellerstein, Herbert M. Sauro

**Affiliations:** Department of Bioengineering, University of Washington, Seattle, WA 98105; eScience Institute, University of Washington, Seattle, WA 98105

**Keywords:** Cascade, Phosphorylation Cycle, First-Order Ultrasensitivity, Frequency Response

## Abstract

Signal transduction from a cell’s surface to cytoplasmic and nuclear targets takes place through a complex network of interconnected pathways. Phosphorylation cycles are common components of many pathways and may take the form of a multi-layered cascade of cycles or incorporate species with multiple phosphorylation sites that effectively create a sequence of cycles with increasing states of phosphorylation. This work focuses on the frequency response and sensitivity of such systems, two properties that have not been thoroughly examined. Starting with a singularly phosphorylated single-cycle system, we compare the sensitivity to perturbation at steady-state across a range of input signal strengths. This is followed by a frequency response analysis focusing on the gain and associated bandwidth. Next, we consider a two-layer cascade of single phosphorylation cycles and focus on how the two cycles interact to produce various effects on the bandwidth and damping properties. Then we consider the (ultra)sensitivity of a doubly phosphorylated system, where we describe in detail first-order ultrasensitivity, a unique property of these systems, which can be blended with zero-order ultrasensitivity to create systems with relatively constant gain over a range of signal input. Finally, we give an in-depth analysis of the sensitivity of an n-phosphorylated system.

## 1 Introduction

Protein signaling pathways communicate information from external signals to both nucleus and cytoplasmic processes in order to modulate cell responses. These pathways engage in various types of signal processing, such as integrating signals over time [34], converting signal strength to signal duration [1], and converting graded signals to switch-like behaviors [7]. In eukaryotes especially, these pathways tend to be highly interconnected, encompassing cross-talk and signal processing between multiple pathways. A large number of signaling pathways exist that include the RTK/RAS/MAP-Kinase pathway, PI3K/Akt signaling, WNT signaling, as well as many others [2].

A common motif found in signaling pathways is the phosphorylation cycle. In these cycles, a protein is phosphorylated in response to a signal and dephosphorylated to return the protein to its original state. Often, such cycles form layers or cascades, where one cycle activates the next. Given the ubiquity of phosphorylation cycles, one might be inclined to consider such cycles as fundamental processing units in biochemical cascades [27]. Many signaling proteins are also phosphorylated at more than one site. For example, MEK and ERK can be doubly phosphorylated. In this case, phosphorylation is processive, meaning that phosphorylation occurs in a strict order, resulting in a two-cycle motif structure (Figure 10). We call these a double cycle and the case with a single phosphorylation, a single cycle (Figure 1). Some signaling proteins are phosphorylated on many sites, though it is still uncertain as to the biological significance of such systems. Makevich et al, [18] showed it was possible for a multi-site phosphorylation cycle to exhibit bistability, and Chickarmane et al [3] showed how oscillations could be obtained from competitive inhibition and multi-site phosphorylation. Finally, Thomson et. al [33] made the intriguing observation that multi-site systems could display many stable and unstable steady-states. Multi-site systems are therefore known to generate a wide variety of dynamic behaviors [22]. In this paper, we also describe first-order ultrasensitivity, which is a unique property among multi-phosphorylated systems.

**Figure 1.**
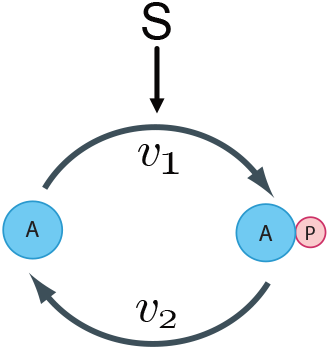
Single phosphorylation cycle with protein A and phosphorylated protein AP. *v*_1_ and *v*_2_ are the reaction rates for the phosphorylation and dephosphorylation reactions, respectively, with signal *S* activating the forward arm, *v*_1_.

The essential steady-state properties of phosphorylation cycles, particularly the single cycle, have been well documented by many authors dating from the late 1970s [31] to the present day [9, 17]. As detailed above, there has also been some interesting work done on multi-site systems. In this article, we explore properties of cascades that have not been studied previously. We look at both single and doubly phosphorylated cycles and focus on their frequency response and sensitivity to perturbations, uncovering some hitherto unrecognized properties.

## 2 Single Phosphorylation Cycle

The single phosphorylation cycle is shown in Figure 1. It involves two proteins, unphosphorylated protein A, and phosphorylated protein AP. Phosphorylation is catalyzed by a kinase, and dephosphorylation by a phosphatase. We assume that the kinase is represented by the signal, S. We examine the properties of the cycle as a function of the signal. This system can be modeled using the following set of differential equations:

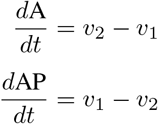

Note that these equations are linearly dependent since either one can be obtained from the other by multiplying by minus one. This is due to mass conservation between A and AP. The moiety, A, is conserved during its transformation to AP and in its conversion from AP to A [11, 25]. Therefore, the total mass of moiety A in the system is fixed and doesn’t change as the system evolves in time. In other words, A + AP = T where T is the fixed total mass of moiety A. This makes the assumption that the synthesis and degradation of protein A and degradation of protein AP is negligible compared to the cycling rate.

Mathematically, the presence of the conservation law means that there is only one independent variable. If we designate the independent variable to be AP, then the dependent variable becomes A and can be computed using a trivial rearrangement of the conservation law: A = T *−* AP where T is the total mass in the cycle.

If we initially assume linear irreversible mass-action kinetics on the forward and reverse limbs, we can write:

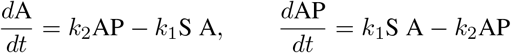

We have assumed, without loss of generality, that the stimulus, S, is a simple linear multiplier into the rate law, *v*_1_. We can readily solve for the steady-state levels of A and AP by setting the independent differential equation to zero, from which we obtain the well-known result:

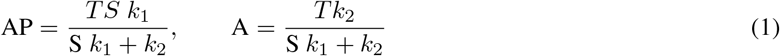

Note that the steady-state level of A is determined from the relation *A* = *T* − AP. The input to the cycle can be modeled by changes to S. We can, therefore, plot the steady-state concentration of AP as a function of the input signal S. This is shown in Figure 2 and illustrates a well-known result.

**Figure 2.**
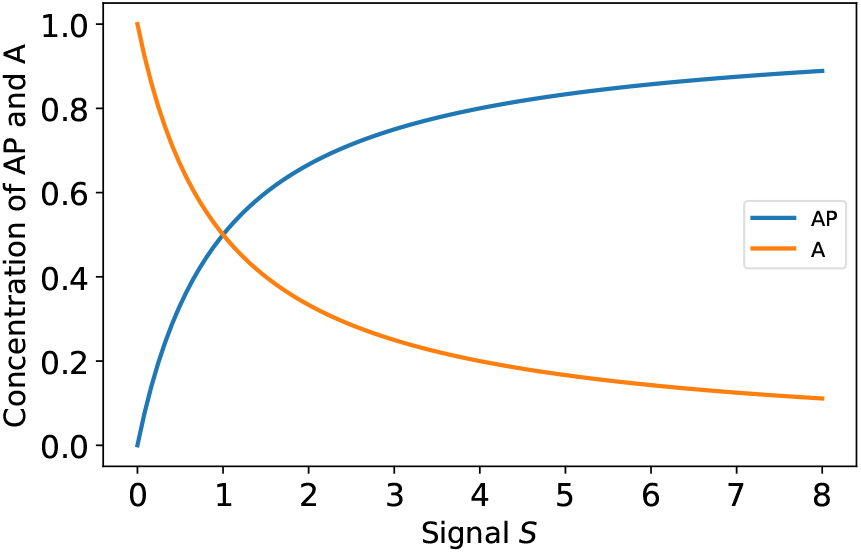
Steady-state plot of AP and A versus signal S for a system with mass-action kinetics. The plot shows the expected and well-known hyperbolic response with no ultrasensitivity.

The response is a rectangular hyperbola (cf. Michaelis-Menten equation). As the stimulus increases, AP increases with a corresponding drop in A due to mass conservation.

### 2.1 Sensitivity Analysis

Of particular interest in this article is to consider the sensitivity of AP to changes in signal S. There are various ways to do this; the most obvious is to evaluate the derivative, *d*AP*/d*S given the steady-state solution to AP. An alternative is to evaluate the scaled derivative, also known as the logarithmic gain [28], since this eliminates units and converts the response into the more intuitive relative change:

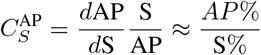

The steady-state equation for the concentration of AP, (1), can be differentiated and scaled to give:

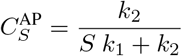

This is a well-known result showing that the sensitivity is always less than or equal to one; that is, a 1% change in S always generates less than a 1% change in AP. Interestingly, the gain of the system is independent of the total mass in the system. Although strictly speaking, these sensitivities are response coefficients [14], because we are assuming the signal elasticity 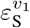 is equal to one, the response and control coefficients are equal to each other. We therefore refer to these sensitivities as control coefficients throughout the article using the letter C.

Much more interesting behavior, but also well-known, is observed if the kinase and phosphatase activity is no longer modeled using simple mass-action kinetics but is modeled using saturable Michaelian rate laws. In this case, the steady-state behavior shows a marked sigmoid response (Figure 3), often termed zero-order ultrasensitivity because the behavior appears when the kinase and/or phosphatase start to operate near the zero-order region of the Michaelian rate law. The degree of sigmoidicity is determined by the degree of saturation of the kinase and phosphatase. This is a well-known result that was shown by Goldeter and Koshland using a detailed mechanistic model [8] of enzyme binding and catalysis. Later Small and Fell [29] showed that zero-order ultrasensitivity could also be demonstrated more generally using small signal sensitivity analysis without having to consider a detailed mechanistic model. This analysis also highlighted the essential properties of a network that were responsible for the zero-order ultrasensitivity. Figure 3 shows the sigmoid response and the scaled and unscaled sensitivities as a function of signal. Of interest is that the peak of the unscaled derivative appears to match the inflection point while the scaled derivative peak is shifted to the left. We don’t have an intuitive explanation for this shift, but we provide proof of its existence in Appendix I.

**Figure 3.**
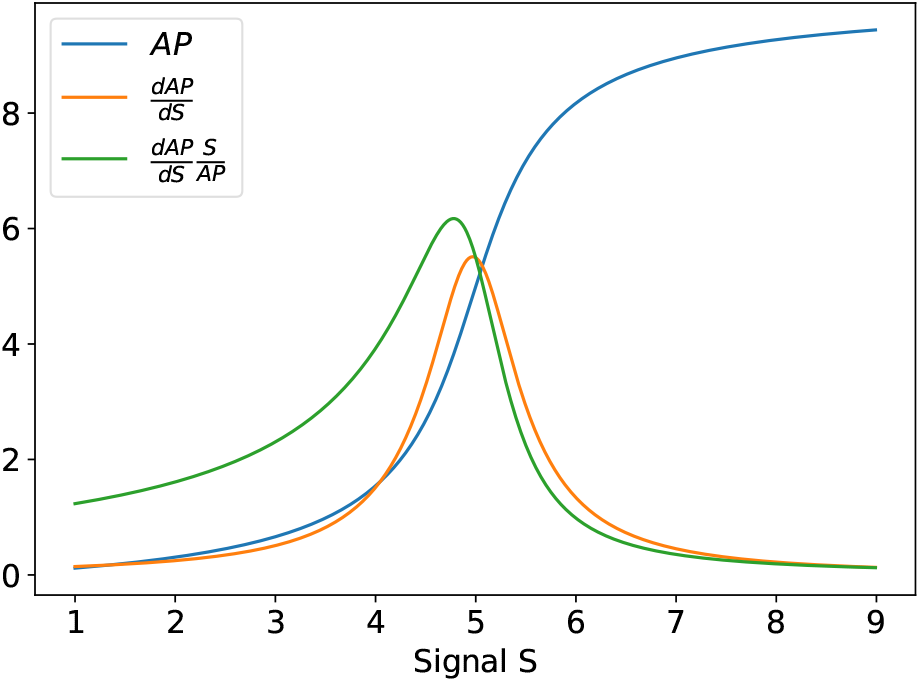
Steady-state plot of AP, the unscaled control coefficient ^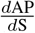^, and the scaled control coefficient 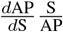over a range of signal, S, values. The kinetics are given by saturable Michaelis-Menten rate laws. Note the sigmoid response in AP and the spike in sensitivity due to the zero-order ultrasensitivity. See Model VIa in the Appendix.

### 2.2 Frequency Response of a Single Cascade

The frequency response describes the steady-state response of a system to sinusoidal inputs at varying frequencies. In general, the sinusoidal inputs are small in amplitude so that even if the system is nonlinear, analytical solutions can be obtained through linearization. Linear systems exhibit two important characteristics in terms of their response to sinusoidal inputs. The first is that the amplitude of the signal can be amplified or attenuated. Secondly, due to inherent delays in the system (for example, the time it takes molecular pools to fill or empty), sinusoidal signals tend to get delayed, resulting in phase shifts. Interestingly, the frequency component of a signal is unaltered when assuming linearity [19]. By examining how a system alters the amplitude and phase of a sinusoidal input, information on the system’s characteristics can be determined. Moreover, a range of sinusoidal frequencies are tested since changes in amplitude and phase are often a function of the input frequency. The result is a particular form of graphical rendering called a Bode plot [13]. These plots invariable come in pairs, one indicating the effect on the amplitude and a second on the phase.

We use the extension of metabolic control analysis to the frequency domain as developed by Ingalls [13] to compute the frequency response. A similar extension was developed by Rao et al [20], which emphasized the application of signal-flow graphs within the context of a frequency response.

The following study includes both analytical analysis and numeric simulations. For the simulations, we assume the model in Appendix VIa (written using the Antimony format [30], which can be readily translated to SBML via the online tool makeSBML https://sys-bio.github.io/makesbml/). We begin by looking at the Jacobian matrix. Due to moiety conservation, the Jacobian only has a single independent element. This can be easily derived using [10]:

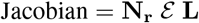

where **N**_**r**_ is the reduced stoichiometry matrix, *ε* the matrix of unscaled elasticites and **L** the link matrix [21]. The unscaled elasticity matrix is a 2 by 2 matrix:

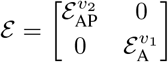

The unscaled elasticity is simply the partial derivative of the reaction rate with respect to a given concentration, hence:

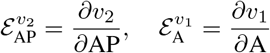

The Link matrix relates the reduced stoichiometry matrix to the full stoichiometry matrix via:

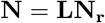

It is relatively easy to manually compute **L** for a single cascade but can be obtained using the function `getLinkMatrix()` in the tellurium package [4]. Likewise, the reduced stoichiometry matrix can be obtained using the function call `getReducedStoichiometryMatrix()`. With this information, the Jacobian can be derived as:

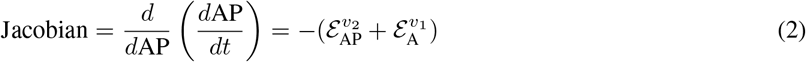

Since the two unscaled elasticities are positive, the Jacobian is negative. Moreover, since the Jacobian is the sum of the two unscaled elasticities, near the steepest portion of the sigmoid curve, these unscaled elasticities are at their minimum. This is what elicits the steep rise in AP but also means that the Jacobian is at a minimum. This is more easily illustrated in a simulation shown in Figure 4. This result may appear to be counter-intuitive since one might expect the sharpest transition in the zero-order ultrasensitive response to be the most responsive and, thereby have the largest value for the Jacobian.

**Figure 4.**
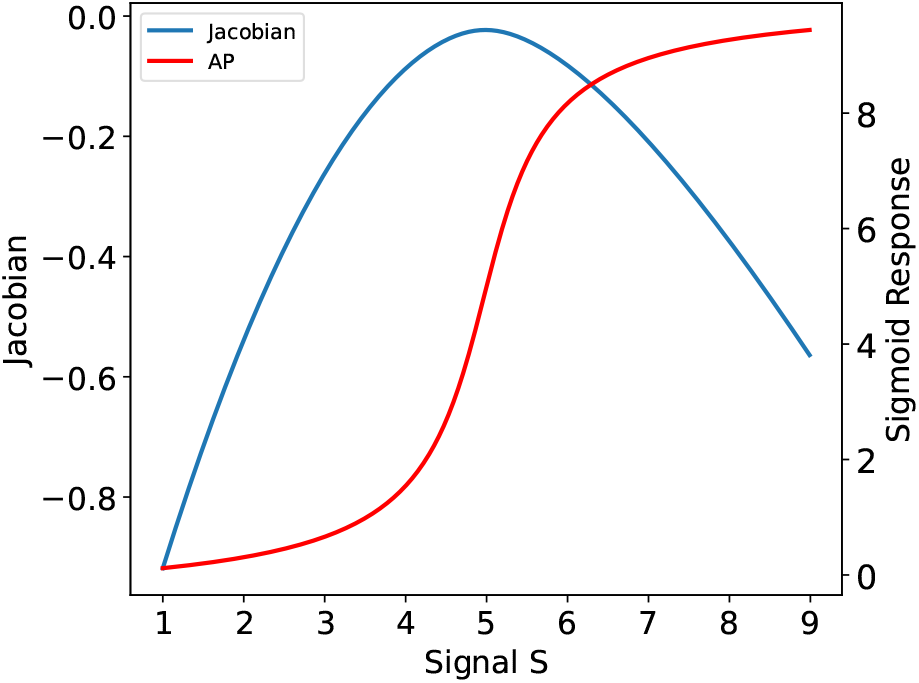
Value of the Jacobian as a function of the input signal. The absolute value of the Jacobian reaches a minimum near or at the steepest point on the sigmoid curve for AP.

The frequency response can be derived analytically using the frequency domain extension by Ingalls [13], Since the system is non-linear this approach necessitates the linearization of the system. The details of the derivation are given in Appendix II. The result is the following transfer function where *s* is a complex variable:

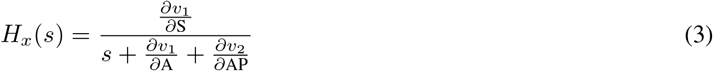

This is a classic first-order system. Its pole is 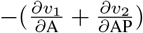 Note that the pole is always negative, and so the system is stable. Further, the speed of response is faster as 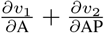increases. Eq. 3 can be written in standard form:

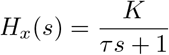

where the gain *K* is given by 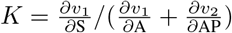 and *τ*, the time constant by:

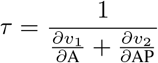

The time constant indicates the responsiveness of the system. The smaller *τ*, the more responsive, and *τ* is smaller if the pole has a larger magnitude. Put differently, the smaller the unscaled elasticities (and hence more zero-order ultrasensitivity), the slower the system is to respond.

The bandwidth (The frequency where the amplitude ratio drops by a factor of 0.707) of a first-order system is simply 1*/τ*. Hence, when the system moves through the steepest portion of the sigmoid curve, its bandwidth is at a minimum. This is also shown in Figure 5, which plots the bandwidth as a function of the signal. This also matches the earlier observation that at the steepest point in the sigmoid curve, the system is least responsive in time. Hence when the system is most responsive to steady-state changes, it is least responsive in time.

**Figure 5.**
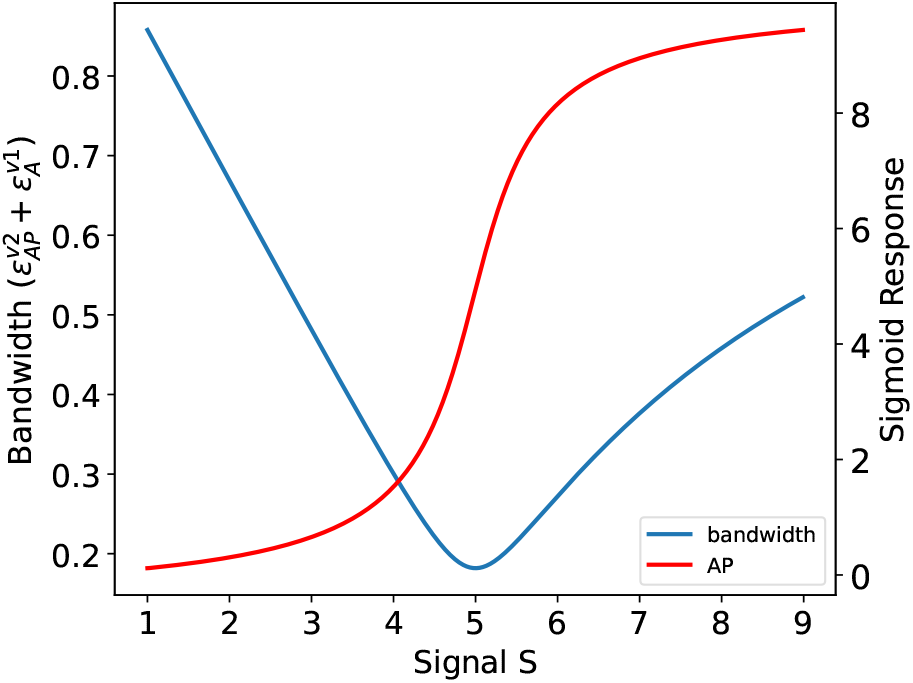
Bandwidth as a function of the input signal for a single cycle. The bandwidth reaches a minimum near or at the steepest point of the sigmoid curve for AP.

The frequency response can be obtained by switching the complex variable, *s*, to the complex frequency *jω* and plotting the Bode plots for the amplitude and phase. These plots are shown in Figure 6 and were computed using the Python Tellurium [4] utility, ‘frequencyResponse’ found at https://github.com/sys-bio/frequencyResponse. This shows a typical response for a low-pass filter. Of more interest is plotting the Bode plots as a function of signal. This results in two 3D plots for amplitude and phase, Figure 7. The amplitude plot clearly shows the reduction in the bandwidth as the signal passes the point of steepest response in the sigmoid curve (At around S=5). At low and high signal levels the bandwidth increases. The work by Gomez-Uribe et. al [9] came to a similar conclusion but by doing simulations on a specific mechanism, and some limited analytical work. Moreover, Thattai and Oudenaarden [32] investigated the effect of zero-order ultrasensitivity on how noise is transmitted and showed attenuation in noise which is consistent with this result.

**Figure 6.**
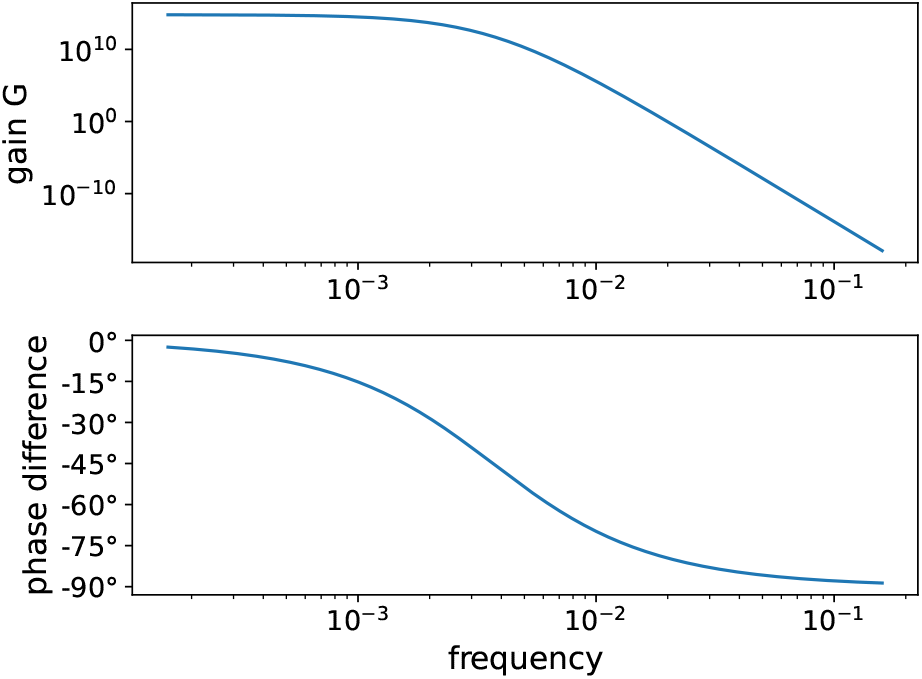
Bode plot for the single cycle system showing the response of AP as a function of signal frequency. The plots show the typical behavior of a low-pass filter with the phase shift reaching a maximum shift of 90 degrees. Both curves were computed when the signal, S, equaled 5.

**Figure 7.**
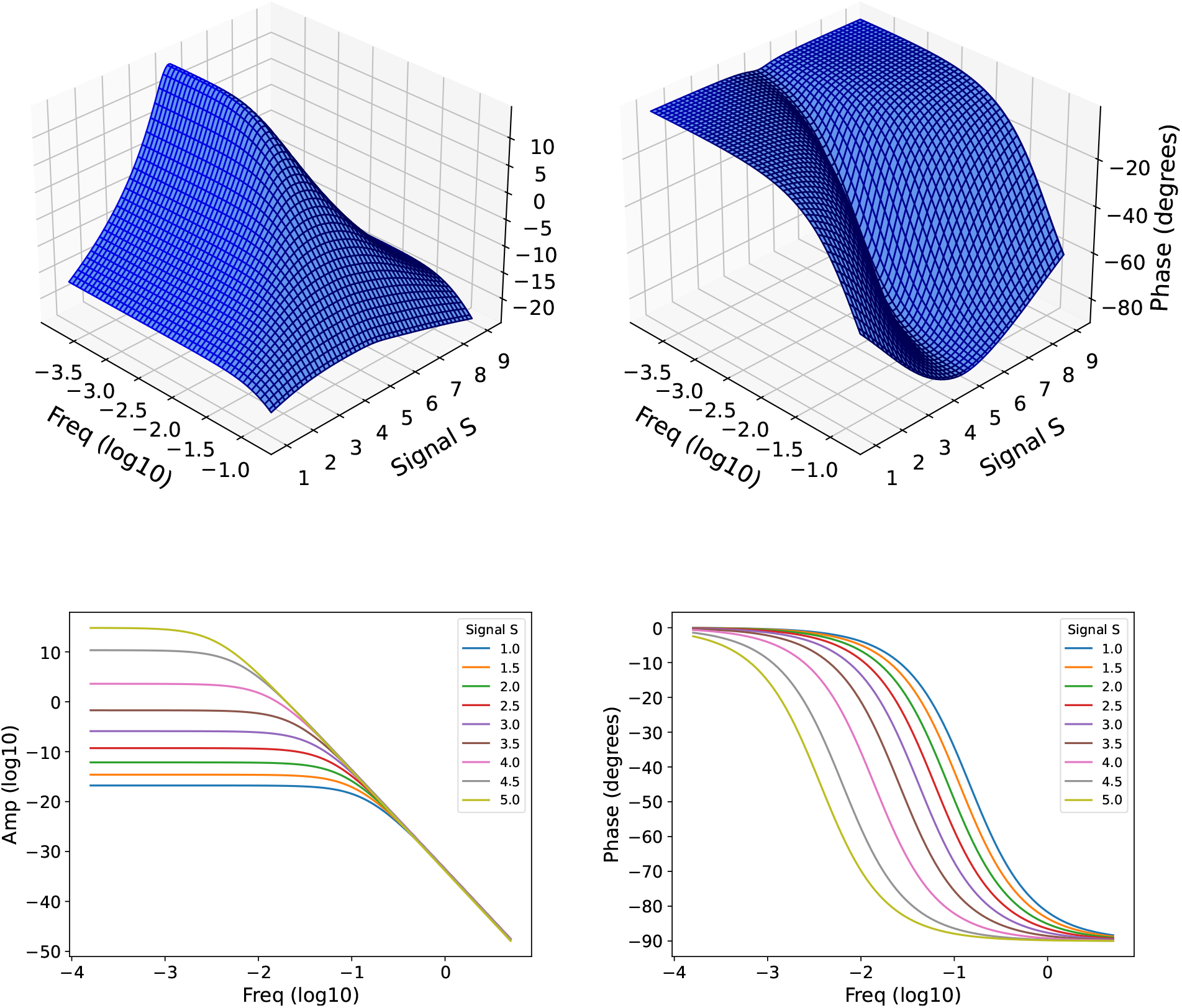
Top: 3D Bode plots for amplitude (left) and phase (right) as functions of frequency and signal S for a single cycle system with output AP. Bottom: 2D slices of the amplitude and phase vs frequency at increasing levels of signal S up to the critical point of the system.

### 2.3 Frequency Response of Two-layered Cascade

We are now going to look at the frequency response for a two-layered single-cycle system. The two-layered model we used is given in Appendix VIb and Figure 8 shows a schematic of the network. The transfer function (and therefore the frequency response) for the two-layered system with respect to BP and the input signal can be derived in a similar manner as before and results unsurprisingly in a second-order system (note there are only two independent variables in this system, AP and BP):

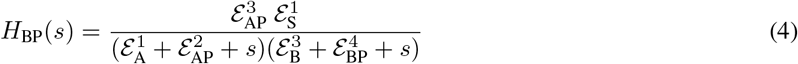

The derivation is given in Appendix III. The transfer function can be converted to the standard second-order form that includes the damping ratio, *ζ* such that *ζ* can be shown to be:

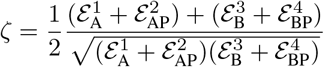

As described in the Appendix III, the value of this expression is always greater than one. This means that all transient behavior of the two-layered cycle is monotonic in nature. The denominator in the transfer function (4) also has two negative real roots again indicting monotonic behavior for the system’s dynamics in response to changes in signal.

**Figure 8.**
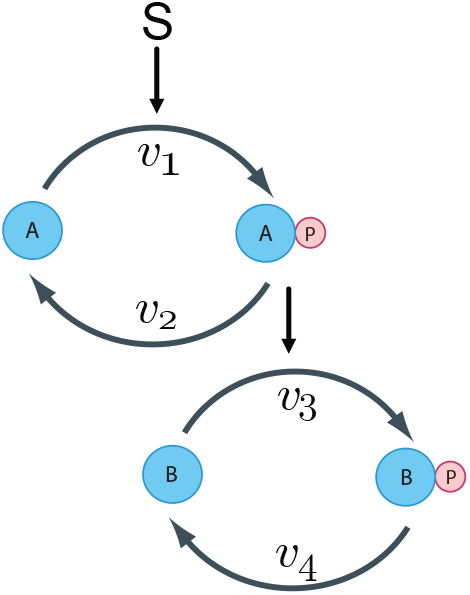
Two single cycles in a layered structure. BP is the output.

The equation for the bandwidth of a second-order system is given by the standard relationship:

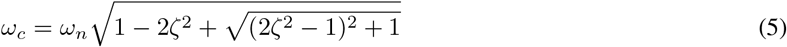

Figure 9 displays the damping ratio and bandwidth for both aligned and unaligned cycles. In the unaligned case (left column), the upstream and downstream cycles reach parity at distinct signal strengths. That is *A* = *AP* and *B* = *BP* occur at different values of S as is evident by the steady-state concentrations for AP and BP in rows 1 and 2, respectively. In the aligned case, the cycles are in parity at the same value of S. This is done by adjusting the parameters so that the *apparent* forward and reverse catalytic constants are equal for each cycle at the same value of S. For the upstream cycle (*A* ⇌ *AP*) that means *k*_1_*S* = *k*_2_ or, for *k*_1_ = 0.5 and *k*_2_ = 0.7 we have *S* = 1.4. For the downstream cycle (*B* ⇌ *BP*) we need *k*_3_*AP* = *k*_4_ when the upstream cycle is balanced (*AP* = 5). Note that there is no retroactivity [6, 16] from the downstream cycle on on AP. Adjusting the parameters so that *k*_4_*/k*_3_ = 5 we have set *k*_3_ = 0.7 and *k*_4_ = 3.5 (one of many possible solutions).

**Figure 9.**
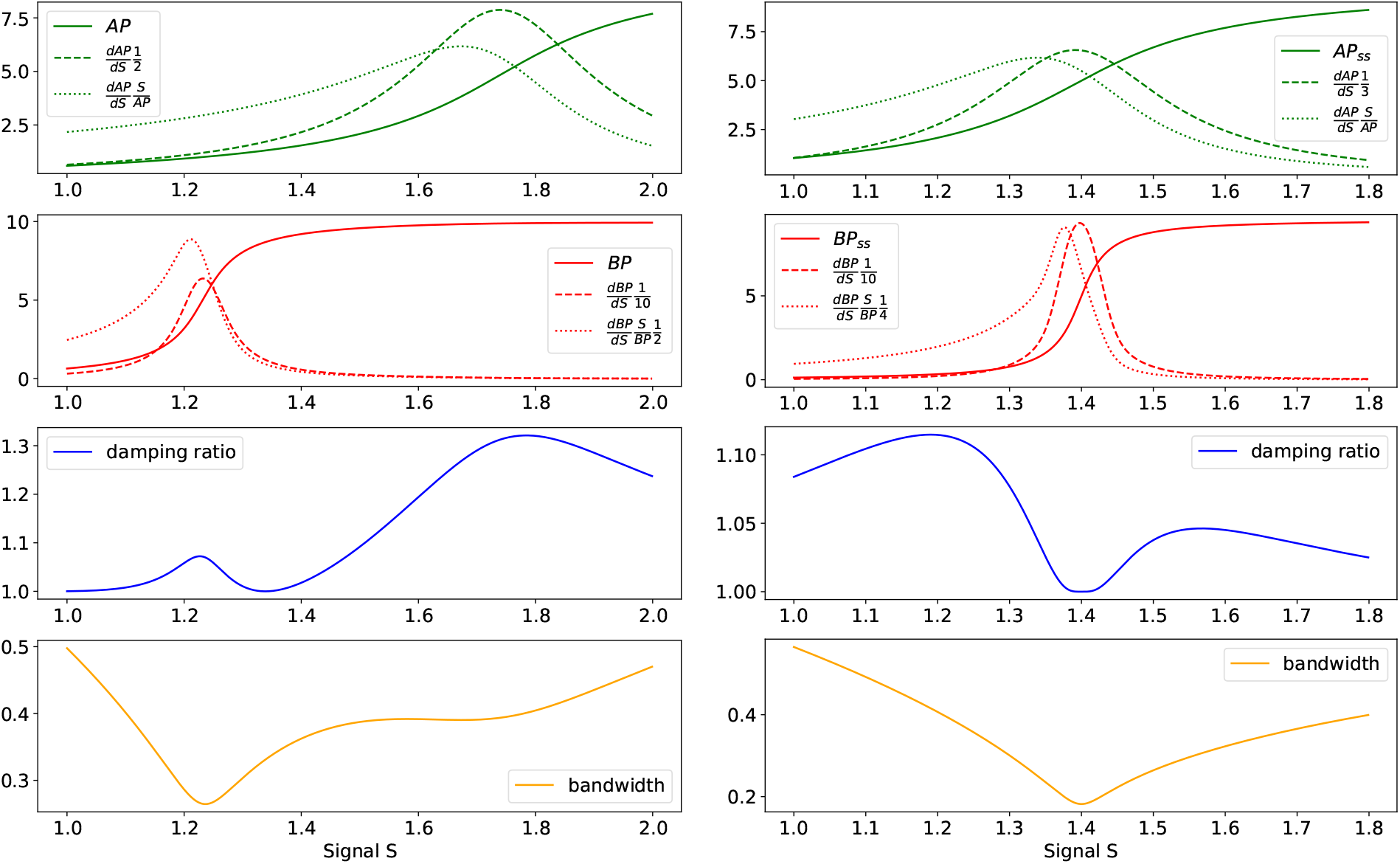
Two layered single cycle system with unaligned (left: *k*_1_ = 0.4, *k*_4_ = 0.7) and aligned (right: *k*_1_ = 0.5, *k*_4_ = 3.5) cycles. Rows 1 and 2 show concentrations and scaled/unscaled control coefficients for AP and BP over changes in signal strength S. Rows 3 and 4 show changes in the damping ratio and bandwidth over changes in S.

**Figure 10.**
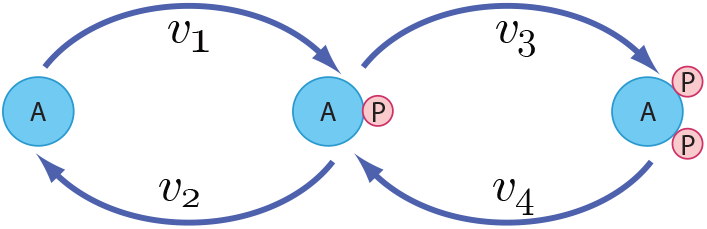
Double Cycle model

As expected, parity for each cycle in the unaligned case is reached at different signal strengths (Figure 9), as does the peaks for the scaled and unscaled control coefficients. For signal strengths ranging from 1 to 2 the damping ratio has two peaks that are near (but not at) the point of parity for the two cycles. For example, the (*A* ⇌ *AP*) cycle reaches parity at a signal signal strength of *S* = 1.75, but the right peak in the damping ratio is closer to *S* = 1.784. This can be explained by considering the components of the damping ratio (Appendix VII Figure 18 left). Note that 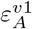 and 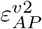 are both relatively low around *S* = 1.75. In fact, the sum 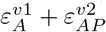 reaches a simulated low of 0.181818 at *S* = 1.75 As this is a multiplicative factor in the denominator, the damping ratio is thus amplified around this point. The same argument can be made for the damping ratio peak near parity for the (*B* ⇌ *BP*) cycle.

The low in the damping ratio for the unaligned system falls between the two points of parity at a simulated signal strength value of 1.339. This is also the point at which sums 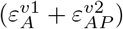 and 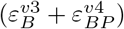 are closest together (within the given range of S). As was shown in Appendix III, the damping ratio can be put in the form of the AM-GM inequality 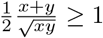 Thus its value is minimized when *x* = *y* or 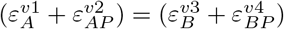.This is true for the aligned case as well, but in that case the sums are equal at points of parity, which are in turn, at the same signal strength (*S* = 1.4) (Appendix VII Figure 18 right). So, instead of a damping ratio peak near the points of parity, as in the unaligned case, we have a minimum. For the bandwidth in the unaligned case there are two minimums, neither of which line up with the points of party, or the damping ratio peaks. However, in the aligned case there is a single minimum that resides at *S* = 1.4, the same signal strength as the damping ratio minimum and the points of parity.

## 3 Double Phosphorylation Cycle

It is very common to find double phosphorylation cycles in protein signaling networks (Figure 10 and Appendix VIc for the antimony model). For example, the MAPK cascade contains two such double cycles. Previous work [12] has reported on some of the zero-order ultrasensitivity properties of such systems. A key feature of double cycles is that they can elicit moderate ultrasensitivity even in the linear regime. However, as we’ll see, the nature of the ultrasensitivity is different from zero-order ultrasensitivity.

We first consider the case when each reaction is governed by simple linear mass-action kinetics. If we assume a stimulus, S, activates *v*_1_ and *v*_3_ we can write the differential equations for this system as:

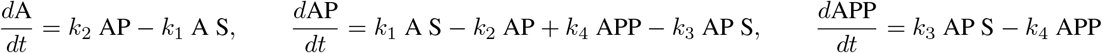

Noting that the total mass of the system is A + AP + APP = T, we can solve for the steady-state to yield:

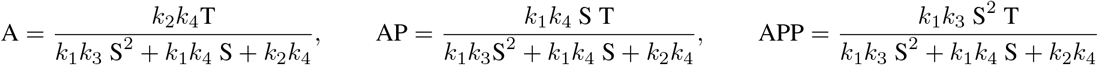

The response of APP to changes in signal can be evaluated by differentiating with respect to signal *S*:

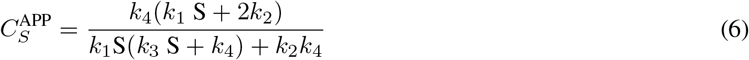

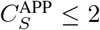 for all positive values of signal strength and parameters *k*_1_ - *k*_4_ (see Appendix VII for a proof). A useful analysis can be obtained by doing a small signal analysis. The proof is given in Appendix V, but the response of APP to changes in the stimulus is given by:

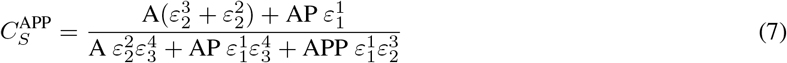

where the 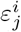 are the scaled elasticity coefficients. Equation (7) looks a little complicated but can be simplified by assuming all reactions are first-order. Under these conditions, all the elasticities equal one so that the equation reduces to something much more manageable:

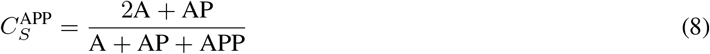

This indicates that given the right ratios for A, AP, and APP, it is possible for 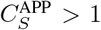. The maximum value the equation can reach is when A and AP are zero, where at this point 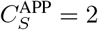. Thus, the maximum is 2.

An important observation is that the character of the response is very different from the more well-known zero-order ultrasensitivity. We can see the difference by looking at the scaled sensitivities shown in Figure 11. On the left, we see the response when the kinetics are linear mass-action. This results in what we call **first-order ultrasensitivity**. It is characterized by the response starting at the maximum value and then decreasing to zero at high signal levels. In contrast, on the right panel, we see the effect of saturable kinetics. However, this is not a purely zero-order ultrasensitivity response. A purely zero-order sensitive response starts at one, spikes close to the inflection point, then rapidly decreases (See Figure 2). For a double cycle, we get a blend of first and zero-order ultrasensitivity. We can see this more clearly in Figure 12. At a moderate level of zero-order ultrasensitivity (At around Km = 6), we achieve a relatively constant gain from the system up to the inflection point. Kms below this result in the appearance of the characteristic spike. This is an interesting behavior that may permit evolution to develop amplifiers with a relatively constant gain without the need for negative feedback [27, 26].

**Figure 11.**
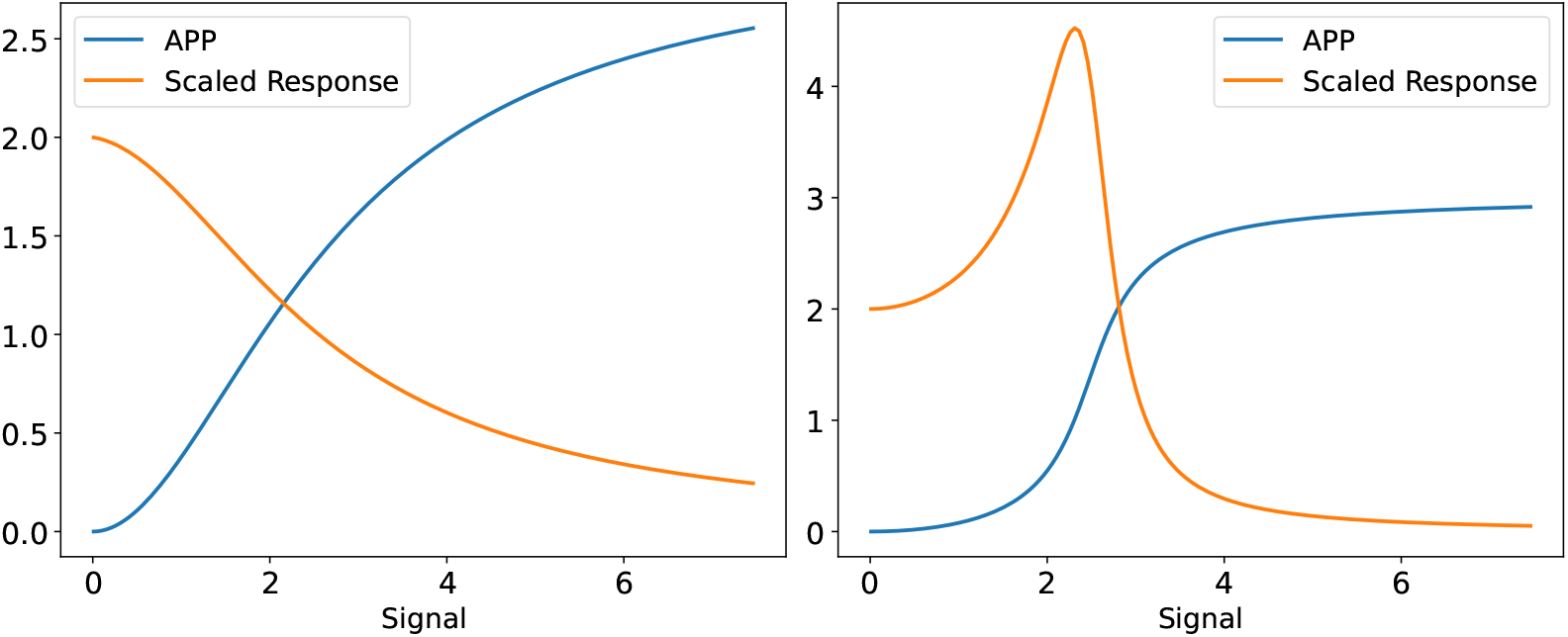
Response of a double cycle to an input signal. Left Panel: Linear mass-action kinetics showing first-order ultrasensitivity; right Panel: Saturable kinetics showing, in this case, a blend of first and zero-order ultrasensitivity. Note how the sensitivity starts at two, not one, as it would in a purely zero-order response (Figure 3.

**Figure 12.**
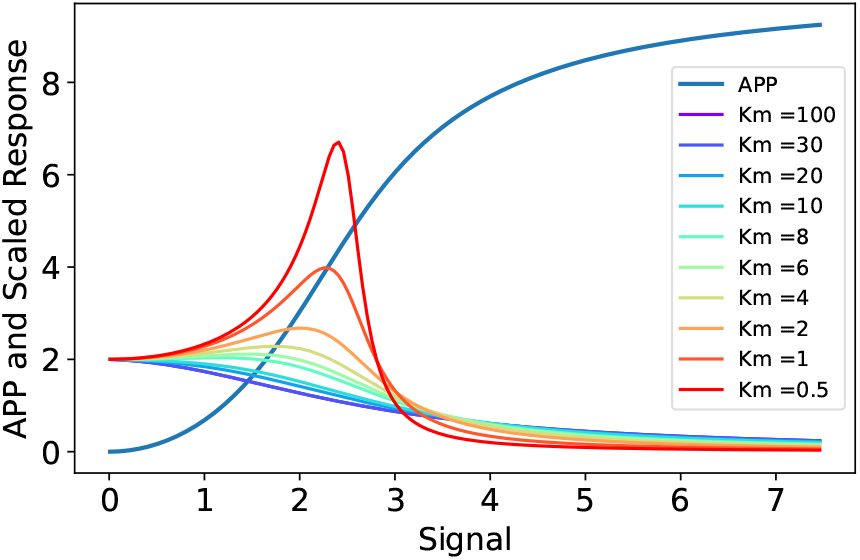
The result of blending first and zero-order responses in a double cycle. High Kms result in pure first-order ultrasensitivity. As the Kms are reduced, more zero-order sensitivity is blended into the first-order ultrasensitive response. Zero-order ultrasensitivity gives us a spike at higher saturation.

One final interesting observation is we see a similar response when comparing the response of a Hill equation to the more complex Monod–Wyman–Changeux model in enzyme kinetics [23]. The response of a Hill equation to the ligand is to first start at the maximum sensitivity and then decrease. This is similar to the first-order ultrasensitivity response. In contrast, the response to a ligand for a Monod–Wyman–Changeux model is to start at 1.0, then peak to a maximum, and then decrease after. This is very similar to the zero-order response.

## 4. Cascades with Many Cycles

This section provides results for single-layer cascades with an arbitrary number of cycles. A cascade has *N* species, and so there are *N−* 1 cycles. In this model, we use *S*_*i*_ to indicate the ith species in the system. Mass is conserved, and so Σ _*n*_ *S*_*n*_ = *T*. We only consider linear mass-action kinetics. Species are organized as in Figure 15, with the subscript of the species increasing from left to right. Odd-numbered kinetic constants refer to reactions that produce a species with a larger subscript. We use mass-action kinetics, for example, *S*_1_ *→ S*_2_ proceeds at the rate *v*_1_ = *k*_1_*S*_1_. Even numbered constants go in the reverse direction. For example, *S*_2_ *→ S*_1_ proceeds at the rate *v*_2_ = *k*_2_*S*_2_.

### 4.1 Steady State Solution With Mass-Action Kinetics

We begin by considering the steady-steady mass or concentration (we use the two interchangeably) of *S*_*n*_ for a cascade with mass-action kinetics. At steady-state, the rate at which mass leaves *S*_*n*_ to the right has to be the same as the rate at which mass enters *S*_*n*_ from the right. Or, *v*_2*n*_*−* _1_ = *v*_2*n*_, and hence *k*_2*n*_ *−*_1_*S*_*n*_ = *k*_2*n*_*S*_*n*+1_, where *k*_*i*_ *>* 0. From this, we infer

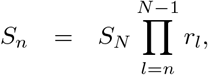

where 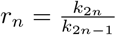. The product of the ratio of the kinetic constants occurs frequently in our analysis, and so we define

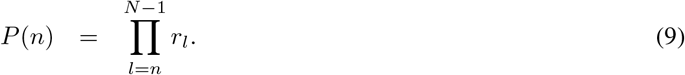

*P* (*n*) is the relative mass of species *S*_*n*_; that is relative to the concentration of *S*_*N*_. Using the constraint that mass is constant,

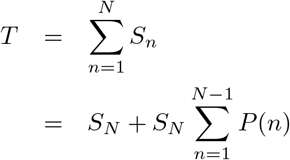

From this, we obtain the steady-state solutions.

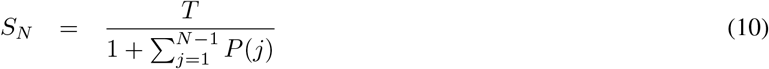

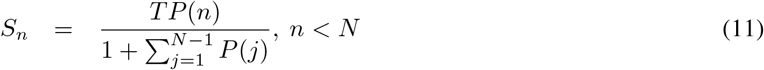

The denominator in these equations is the total relative mass. The numerator is the relative mass for a particular species multiplied by the total mass.

One insight from the foregoing is that the steady does not depend on the absolute value of the kinetic constants, just the ratios *r*_*n*_. Also, note that if for 1*≤ l ≤N−* 1 we have *r*_*l*_ *<* 1, then *P* (*n*) *< P* (*n* + 1), and so at steady-state more mass is associated with species closer to *S*_*N*_. If *r*_*l*_ *>* 1, then more mass is associated with species closer to *S*_1_. Last, consider the case where there is an *n*^*⋆*^ such that (a) *r*_*n*_ *<* 1 for *n < n*^*⋆*^ and (b) *r >* 1 for *n ≥n*^*⋆*^. Then, *P* (*n*^*⋆*^) = *max*_*n*_*P* (*n*) *>* 1, and so *S*_*n*\*_ has the most mass of the species.

### 4.2 Control Coefficients

There are two ways to derive the control coefficient relationships. The first approach is we can assume linear mass-action kinetics for all steps and solve for the steady-state concentrations as we did in the last section. We can then derive the scaled derivatives from the analytical solutions. The second approach is to assume nonlinear kinetics and linearize the system and express the response in terms of the system elasticities. We describe both approaches.

#### 4.2.1 Responses based on mass-action kinetics

This section derives control coefficients with a focus on the last stage in the cascade, *S*_*N*_. Control coefficients are calculated relative to the ratios *r*_*m*_; that is, we have a signal that only affects the ratio *r*_*m*_ associated with *S*_*m*_. From this, we calculate 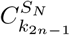 and 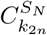.

Consider 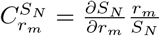. Note that

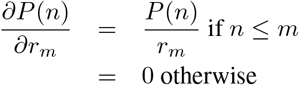

and so

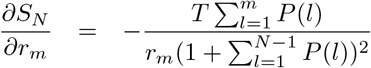

We now calculate the control coefficients as

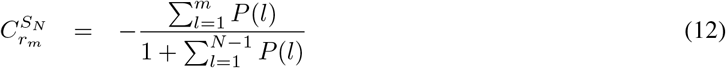

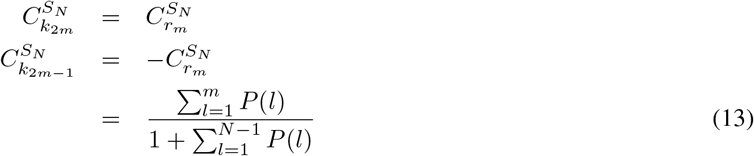

Eq. 13 provides an interesting insight. Control of *S*_*N*_ by *S*_*m*_ is possible only if we can transfer mass between *S*_*N*_ and *S*_*m*_. In general, we want *S*_*N*_ to be small so that little mass needs to be transferred to achieve greater control. So for *S*_*m*_ to control *S*_*N*_, either *S*_*m*_ must be large or *S*_*m*_ must mediate mass transfers from *S*_*n*_, *n < m*. That is, the control coefficient 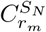 is large if the sum of the mass of *S*_*n*_, *n ≤ m* is large. This summation is the numerator of Eq. 13. The denominator is a normalization constant.

We can use Eq. 10 and Eq. 13 to design a cascade to control species *S*_*N*_. By design, we mean specifying values of the *r*_*m*_. The primary objective of the design is to provide effective control by making the control coefficients 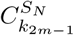as large as possible. A secondary consideration in the design is determining the fraction of mass for each species since there may be constraints related to species concentrations.

Our first observation from Eq. 13 is that 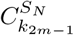 has a maximum value of 1. This is apparent since the summation in the numerator is part of the summation in the denominator. We maximize 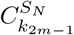by making the numerator of the summation very large. Note that *P* (*l*) *>* 0 and there are more terms in the numerator summation for larger *m*. So, control coefficients are monotonically increasing in *m*. That is, 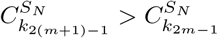

From the foregoing, we can make 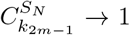 by having 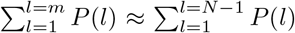. We can make *all* of these control coefficients large if 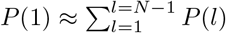. *P*(1) is large if *k*_1_ is small and/or *k*_2_ is large.

Fig. 13 displays the design of a cascade in which *P* (1) *>> P* (*l*) for *l >* 1. Note that by simultaneously controlling all cycles in the cascade, we achieve a total control of 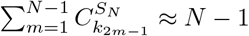. It is interesting to note that the behavior the system exhibits when activating one cycle is very similar to how a linear metabolic pathway responds. While a metabolic pathway relies on product inhibition to transmit changes [24], a series of protein cycles uses movement in a fixed amount of mass to elicit transmission changes. One major difference is that the sum of control coefficients in a metabolic pathway is one, while in the sequence of cycles, the maximum value is equal to the number of cycles. Here we that simultaneously manipulating the reaction rates for the *N −*1 cycles results in a control coefficient of *N−* 1. Since *P* (1) *>> P* (*l*) for *l >* 1, we know that most of the mass in the cascade is *S*_1_.

**Figure 13.**
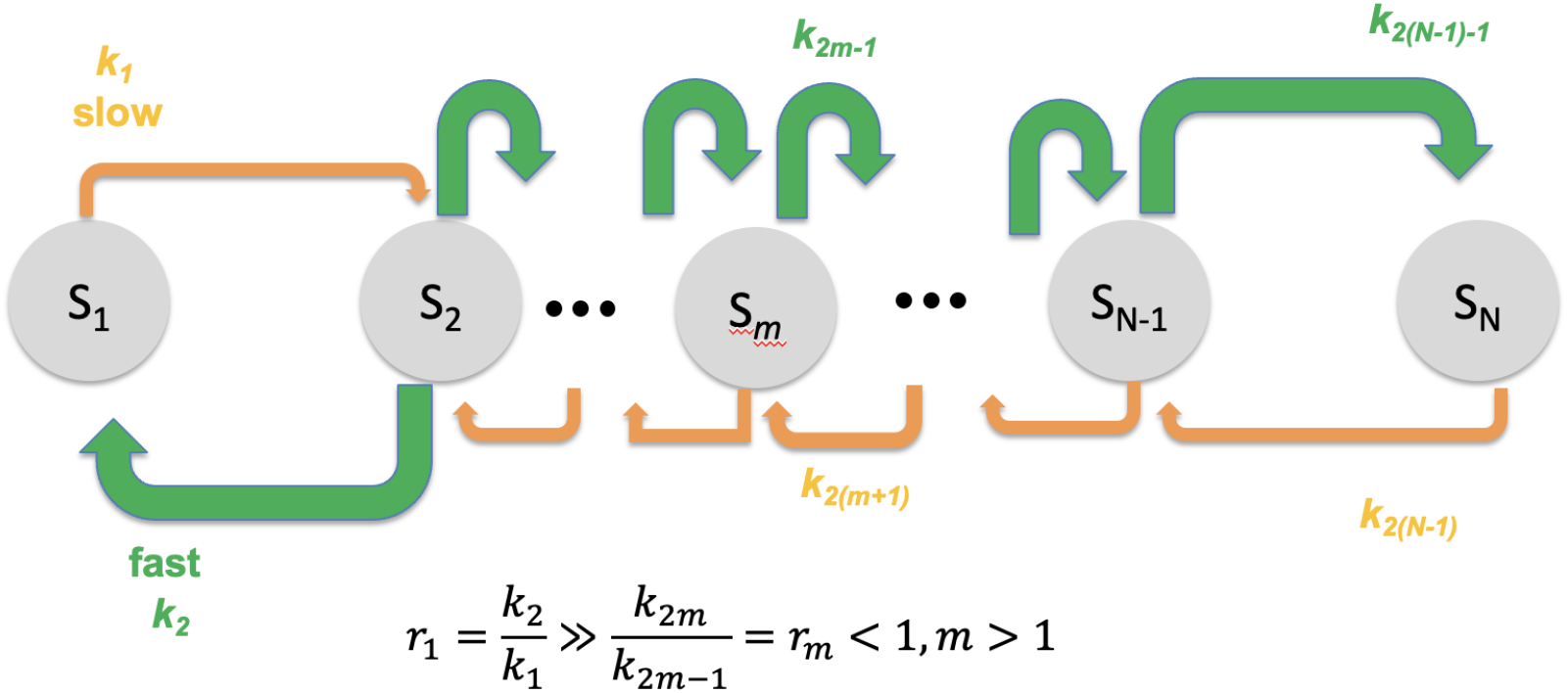
Designing a controllable cascade for changing *S*_*N*_. The cascade has large values of control coefficients for each 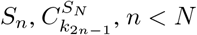, *n < N*. The key to the design is that *S*_1_ has most of the mass so that *k*_2*n−*1_ mediates the transfer of mass between *S*_1_ and *S*_*N*_.

Figure 14 displays control coefficients obtained from Tellurium simulations of a 4 cycle (5 species) cascade (See Appendix VId). The plot in the upper left displays the control coefficients as we vary the value of *r*_1_ for which the control coefficients are calculated. A small value of *r*_1_ results in a small 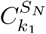. The mass is equally distributed among the other species, and so species closer to *S*_*N*_ mediate the transfer of more mass and hence have larger control coefficients. A large value of *r*_1_ results in most of the mass being *S*_1_, a situation that provides more control when adjusting the kinetic constants. Indeed, a very large *r*_1_ results in the control coefficients converging to 1.

**Figure 14.**
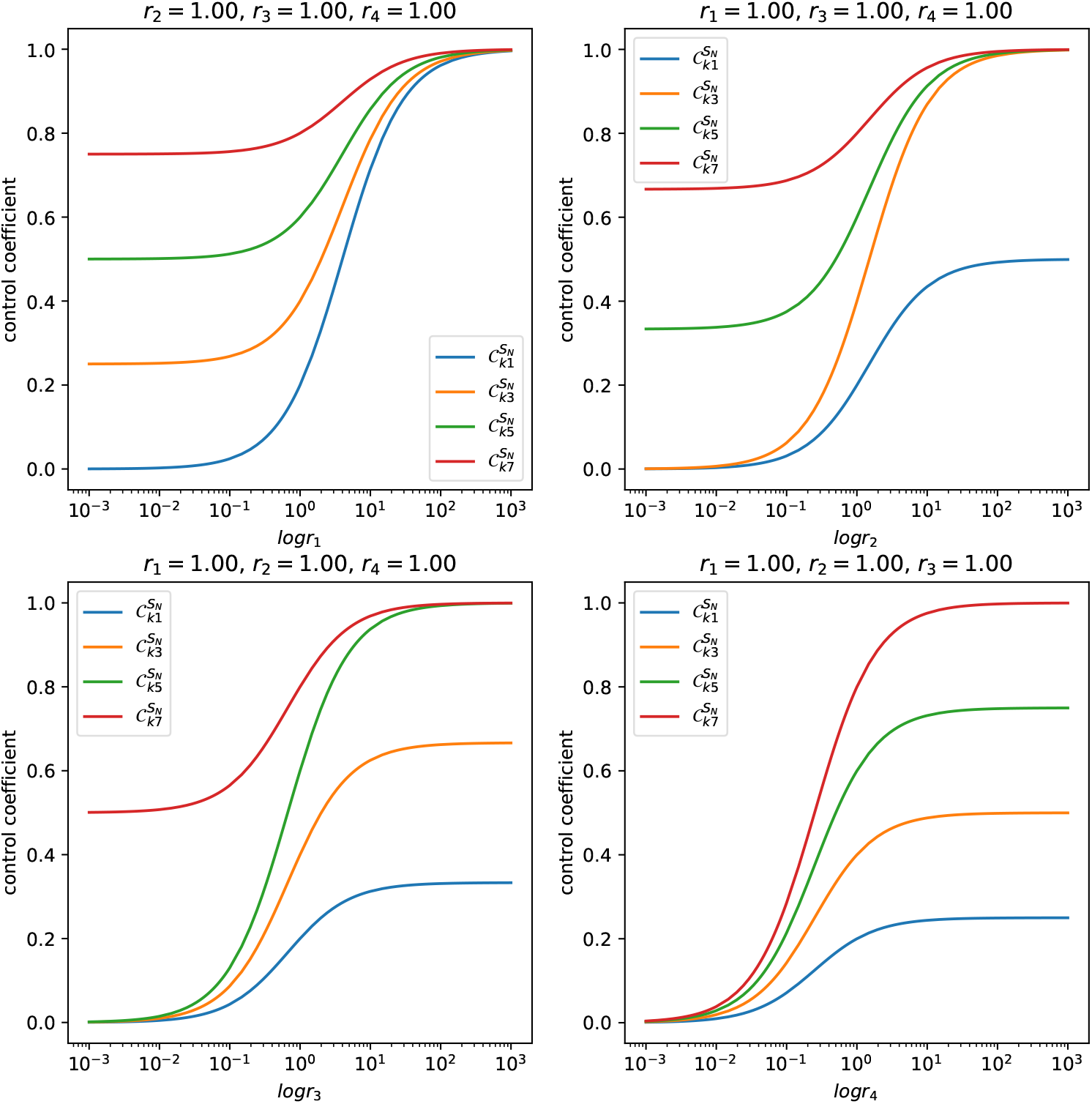
Control coefficients for *S*_5_ in a 5 species cascade. Better control is achieved by having *r*_1_ large (e.g., by making *k*_1_ small). A large *r*_1_ means that most mass is *S*_1_ and so that *S*_1_-*S*_4_ mediate the transfer of mass between *S*_1_ and *S*_5_.

**Figure 15.**
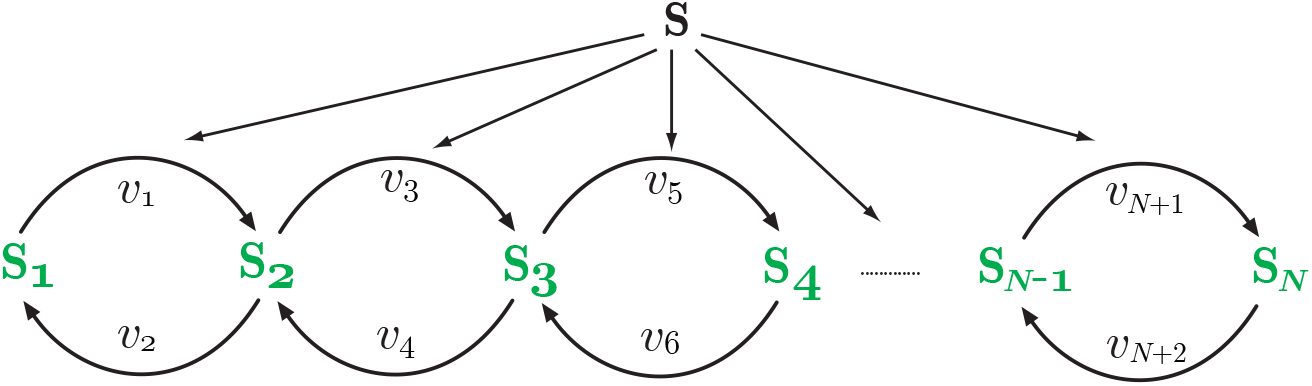
Multiple cycles with S as the stimulus signal.

The other plots explore the effect of varying *r*_*m*_, ∈ *m* {2, 3, 4}. Large values of *r*_2_ result in *S*_2_ having most of the mass. Since *S*_1_ lies to the left of *S*_2_, adjusting *k*_1_ transfers little mass to *S*_*N*_. As a result, 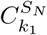 is much smaller in the upper right plot than in the upper left plot. Similar effects can be seen in the bottom row of plots.

#### 4.2.2 Responses based on linearization

In this section, we derive the control coefficients in terms of elasticities and be based on the model shown in Figure 15.

The proof can be found in Appendix V, which illustrates the case for a two-cycle system but the proof can easily be generalized to multiple cycles. For a three-cycle system, where APPP is the output, the response when the signal S activates each forward arm is given by:

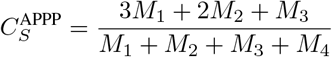

The maximum value that 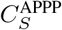 can reach in this case is three. This can be generalized to a system with *N* – 1 cycles (or *N* proteins) where it can be shown that the maximum response is *N* – 1. For example, a system that has six cycles displays a maximum response of six.

Figure 16 shows a plot of the response of a six-cycle system that uses mass-action kinetics for each step.

**Figure 16.**
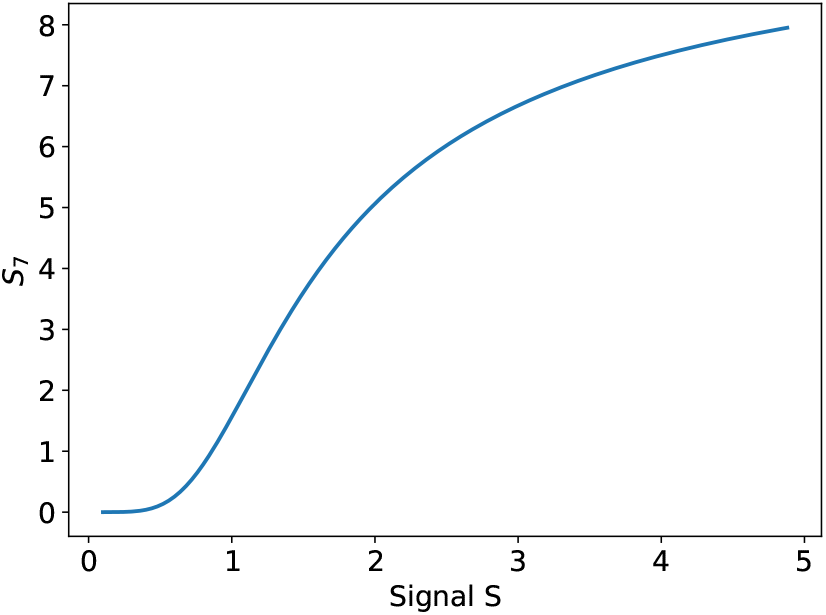
Multiple cycles with S as the stimulus signal.

## 5. Conclusion

In this paper, we discuss two aspects of phosphorylation cycles that have not received as much attention in the past. Specifically, we examine some of the frequency response and sensitivity characteristics of single, double, and multiple-cycle systems. We note that a single cycle behaves as a classic low-pass filter. More interestingly, when operating under zero-order conditions, the bandwidth of the system is a minimum at the most sensitive point on the sigmoid ultrasensitivity curve. This indicates that the system acts as a noise filter in this region of the response. When cascading two single cycles, the system acts as a second-order system. we show that the damping ratio for such a system cannot go below one. This means that all transient dynamics as a result of perturbations to the input signal are always monotonic.

We also examined double cycle sensitivities. Double cycles can show ultrasensitivity even when operating in the sub-saturating regimes of the kinases. We show that a double cycle under these conditions can have a maximum response sensitivity of 2. These call this effect first-order ultrasensitivity to contrast it with the more well-known zero-order sensitivity. We show that first-order ultrasensitivity has unique response properties where its sensitivity is maximum at zero signal, then decreases to zero at high signal levels. There is also an accompanying characteristic plateau near the start before the response starts to decline. Of interest is that as saturation is introduced into the system, zero-order ultrasensitivity emerges but is blended in with the first-order ultrasensitivity. This allows a unique behavior not found in single cycles. Whereas in a single cycle zero-order ultrasensitivity peaks near the steepest portion, then declines rapidly, a blend response allows the sensitivity to remain relatively constant up to the steepest portion. This may have evolutionary significance as it allows the system to have a wider range of constant gain. It is of interest to note that the bottom two layers of the MAPK cascade use double cycles.

Finally, we look at the sensitivity of multiple cycles and note that under first-order ultrasensitivity conditions, the maximum sensitivity is equal to the number of cycles. We also note that the response to changes at individual cycle points can be explained in a similar manner to how control coefficients are distributed in a linear metabolic pathway.

There are some areas we have not considered in this paper due to time constraints. The first is how the frequency response of a single cycle and double cycle compare? Some initial investigations suggest that very little difference exists and a double cycle behaves as an over-damped system. What we do not know is how the frequency response compares under first and zero-order conditions.

## 6. Availability: Software and Models

All Python scripts used to generate the figures and simulations can be found at: https://github.com/sys-bio/frequency_response_paper

## Acknowledgments

We with to thank Steve Wiley and Song Feng for their useful discussions. The research reported in this article was supported by the National Institute of Cancer of the National Institutes of Health under award number U01CA227544. This work was supported in part by the Washington Research Foundation and by a Data Science Environments project award from the Gordon and Betty Moore Foundation (Award #2013-10-29) and the Alfred P. Sloan Foundation (Award #3835) to the University of Washington eScience Institute.

## 7 Appendix

### I Proof for the shift in the scaled sensitivity

Simulations indicate that the scaled response 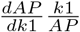peaks just before the maximum slope of the sigmoid plot. The following provides proof of that.

Let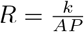 Then at steady-state, we have

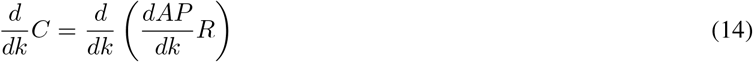

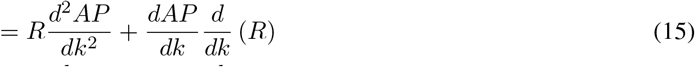

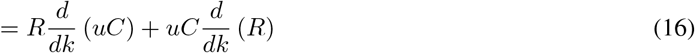

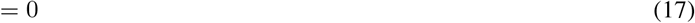

We note there is an extra term in the derivative of the control coefficient with respect to signal, k, that accounts for the *k* dependent scaling term^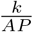^. Thus, there is a shift in the peak for the control coefficient relative to the unscaled coefficient.

### II. Derivation of the frequency response for a single cycle (equation 3)

The basis for the derivation is ([13]):

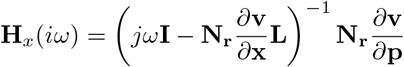

where **N**_**r**_ is the reduced stoichiometry matrix, **L** the link matrix, *∂***v***/∂***x** the unscaled species elasticity matrix and *∂***v***/∂***p** the parameter elasticity matrix.

Using Tellurium, the following was obtained:

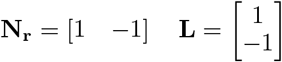

The independent species is AP; hence, the single row in the stoichiometry matrix corresponds to AP. The two columns correspond to *v*_1_ and *v*_2_. The unscaled elasticity matrix is a 2 by 2 matrix given by:

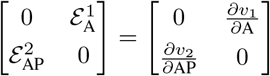

Two entries are zero because we assume no product inhibition on the kinase (*v*_1_) phosphatase (*v*_2_) by AP and A, respectively. The parameter unscaled elasticity matrix is given by:

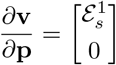

The parameter elasticity matrix only has a single non-zero term because we assume that the only interaction is by signal, S, on *v*_1_. Insertion of these terms into the frequency response expression leads to:

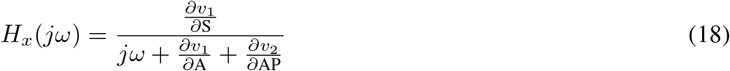

The fact that the system is first-order makes this a simple derivation.

### III. Derivation of the frequency response for two-layers of single cycles (equation 4)

Tellurium was used to derive the reduced stoichiometry (**N**_**r**_) and link matrix (**L**):

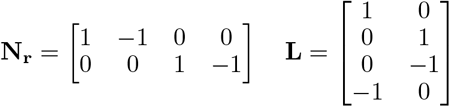

The order of the species in the model was set to ensure that the top two rows of the stoichiometry matrix were AP and BP, respectively. This resulted in the two independent species being AP and BP. As a result, the unscaled elasticity matrix was:

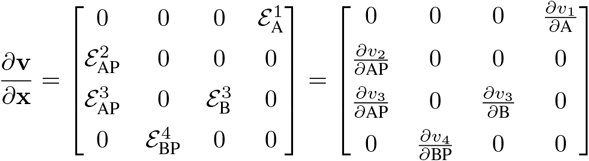

The parameter elasticity is the same as for the single layer except the number of rows is extended to four:

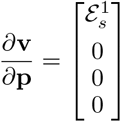

The entry 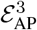 represents the elasticity that connects the two layers together. As before, the frequency response can be derived by inserting these terms into the equation:

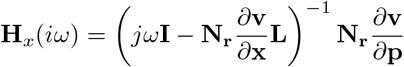

This leads to a second-order system:

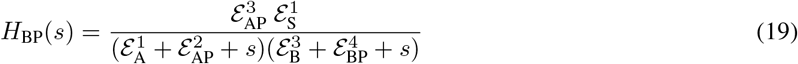

When the denominator is expanded this gives:

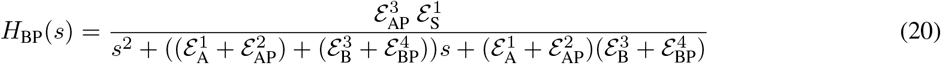

This can be converted into the standard form for a second-order system by dividing top and bottom by 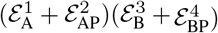, and setting 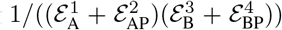to 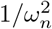, such that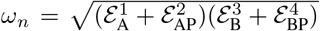. This allows us to rewrite the transfer function as:

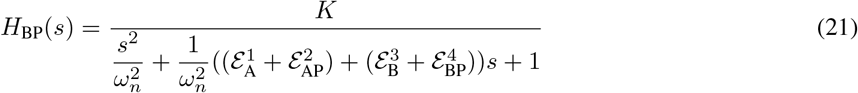

where *K* is the system gain. Finally, we define:

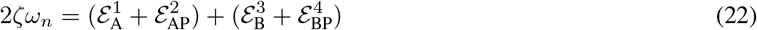

and multiplying top and bottom by 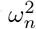, results in:

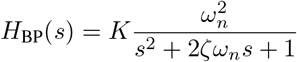

The transfer function is now in the standard form where *ζ* is the damping ratio. Equation 22 can be rearranged to give:

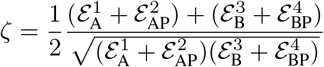

This equation is of the form:

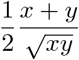

which can be shown to have a value great than one as follows.

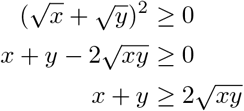

Hence:

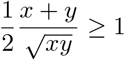

This means that the damping ratio, *ζ* is greater than one. Second-order systems with a damping ratio greater than one cannot admit any damped periodic behavior. This means all dynamic behavior of the two-cycle system must be monotonic.

### IV. Proof for equation 7

Consider the double cycle model shown in Figure 17.

**Figure 17.**
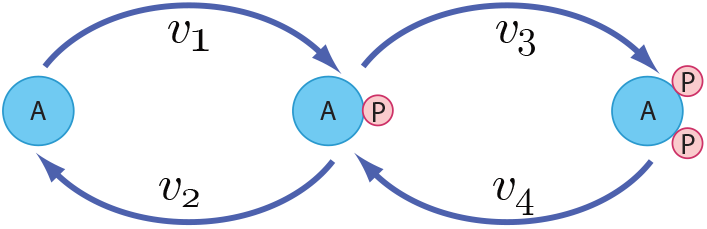
Double Cycle model

To keep things simple, we assume no product inhibition or reversibility in the cycle reactions. Although we are doing a manual derivation for the expression, it is possible to use PyscesToolbox [5], which is an extremely effective tool for deriving control coefficient expressions symbolically and we highly recommend it use it in such cases. However, the manual derivation illustrates the deductive approach that can be used to derive sensitivities within the framework of metabolic control analysis.

The strategy is to first find the expressions for 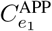 and 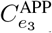 The response to a signal S, is the sum [21, 15] of these expressions assuming the elasticity of S to *v*_1_ and *v*_3_ is one. This is a reasonable assumption since S is usually a kinase, and catalysis is normally first-order. We show the derivation for 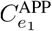. The derivation for 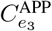 can be done in a similar manner.

To derive 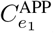, perturb *e*_1_ by an amount *δe*_1_. This changes the steady state from which the following local equations can be obtained (Note that subscripts on the elasticities, 1 = A, 2 = AP, and 3 = APP):

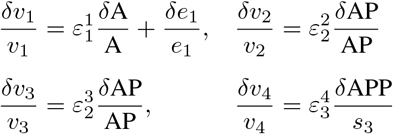

At steady state *v*_1_ = *v*_2_ and *v*_3_ = *v*_4_, though it is **not** necessarily the case that *v*_1_ = *v*_3_. This means that when the steady state changes *δv*_1_ = *δv*_2_ and *δv*_3_ = *δv*_4_. In relative terms we state that: *δv*_1_*/v*_1_ = *δv*_2_*/v*_2_ and *δv*_3_*/v*_3_ = *δv*_4_*/v*_4_. By equating the local equations *δv*_1_*/v*_1_ and *δv*_2_*/v*_2_ we obtain:

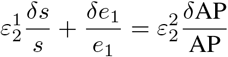

Both sides of the equation can be divided by *δe*_1_*/e*_1_ to give:

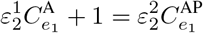

A similar equation can be derived for 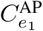 and 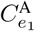 using the *v*_3_, *v*_4_ pair of local equations. In this case, the result is simpler:

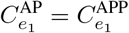

As a result we have two equations and three unknowns 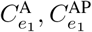, and 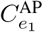. To solve for the three unknowns, a third equation is necessary. The double cycle has a single conservation equation, A + AP + APP = T. Perturbing *e*_1_ by *δe*_1_ does not disturb the total T but changes the distribution of species such that the change in species must be constrained by *δ*A + *δ*AP + *δ*APP = 0. Scaling each term:

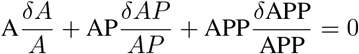

and dividing throughout by *δe*_1_*/e*_1_ yields:

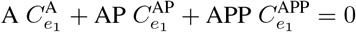

We now have three equations in three unknowns, which can be solved. For example, solving for 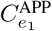 gives:

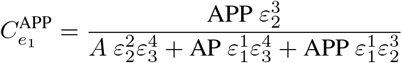

Using the same technique, a solution to 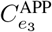 can also be found as follows:

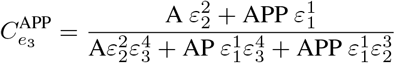

The influence of an external signal, S, is the sum of its interactions therefore 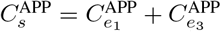 where we assume that the elasticity of the signal on *v*_1_ and *v*_3_ is one. This gives us the total response of *s*_3_ due to changes in the signal S. The sum is given by the equation (23):

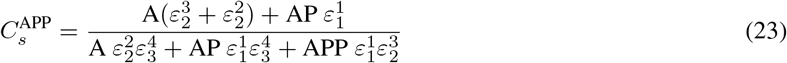

This is equation 7 in teh main text.

### V. Antimony models

#### Va. Single cycle

~~~
 # Declaring AP first ensures that the first row
 # of the stoichiometry matrix will be AP
 # This forces the independentt variable to be AP
 species AP, P
 A -> AP; k1*S*A/(Km + A)
 AP -> A; k2*AP/(Km + AP)
 k1 = 0.14; k2 = 0.7
 A = 10; S = 1; Km = 0.5
~~~

#### Vb. Single cycle two layers

~~~
 species AP, BP, A, B
 v1: A -> AP; k1*S*A/(Km + A)
 v2: AP -> A; k2*AP/(Km + AP)
 v3: B -> BP; k3*AP*B/(Km + B)
 v4: BP -> B; k3*BP/(Km + BP)
 k1 = 0.4 # 0.5
 k2 = 0.7; k3 = 0.7
 k4 = 0.7 # 3.5
 Km = 0.5
 A = 10; B = 10; S = 1
~~~

#### Vc. Double cycle

~~~
 species APP, AP, A
 J1: A -> AP; k1*S*A/(Km + A)
 J2: AP -> A; k2*AP/(Km + AP)
 J3: AP -> APP; k3*S*AP/(Km + AP)
 J4: APP -> AP; k4*APP/(Km + APP)
 k1 = 0.14; k2 = 0.7
 k3 = 0.7; k4 = 0.7; Km = 0.5
 A = 10; S = 1
~~~

#### Vd. N cycle model

~~~
 J1f: S1 -> S2; S1*k1;
 J1b: S2 -> S1; S2*k2;
 J2f: S2 -> S3; S2*k3;
 J2b: S3 -> S2; S3*k4;
 J3f: S3 -> S4; S3*k5;
 J3b: S4 -> S3; S4*k6;
 J4f: S4 -> S5; S4*k7;
 J4b: S5 -> S4; S5*k8;
 k1 = 1; k2 = 1.0;
 k3 = 1; k4 = 1.0;
 k5 = 1; k6 = 1.0;
 k7 = 1; k8 = 1.0;
 S1 = 100; S2 = 0;
 S3 = 0; S4 = 0;
 S5 = 0;
~~~

### VI. Proof that 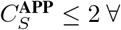 positive values of S and parameters *k*_1_ through *k*_4_ (equation (6))

Proof: Suppose *∃* a positive signal strength S and positive parameters *k*_1_ - *k*_4_ such that

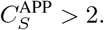

Then

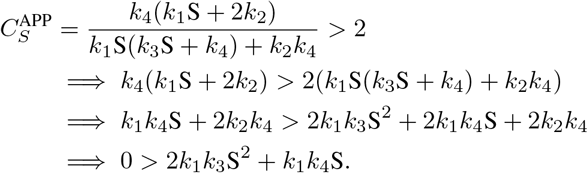

A contradiction. □

### VI. Figures

**Figure 18.**
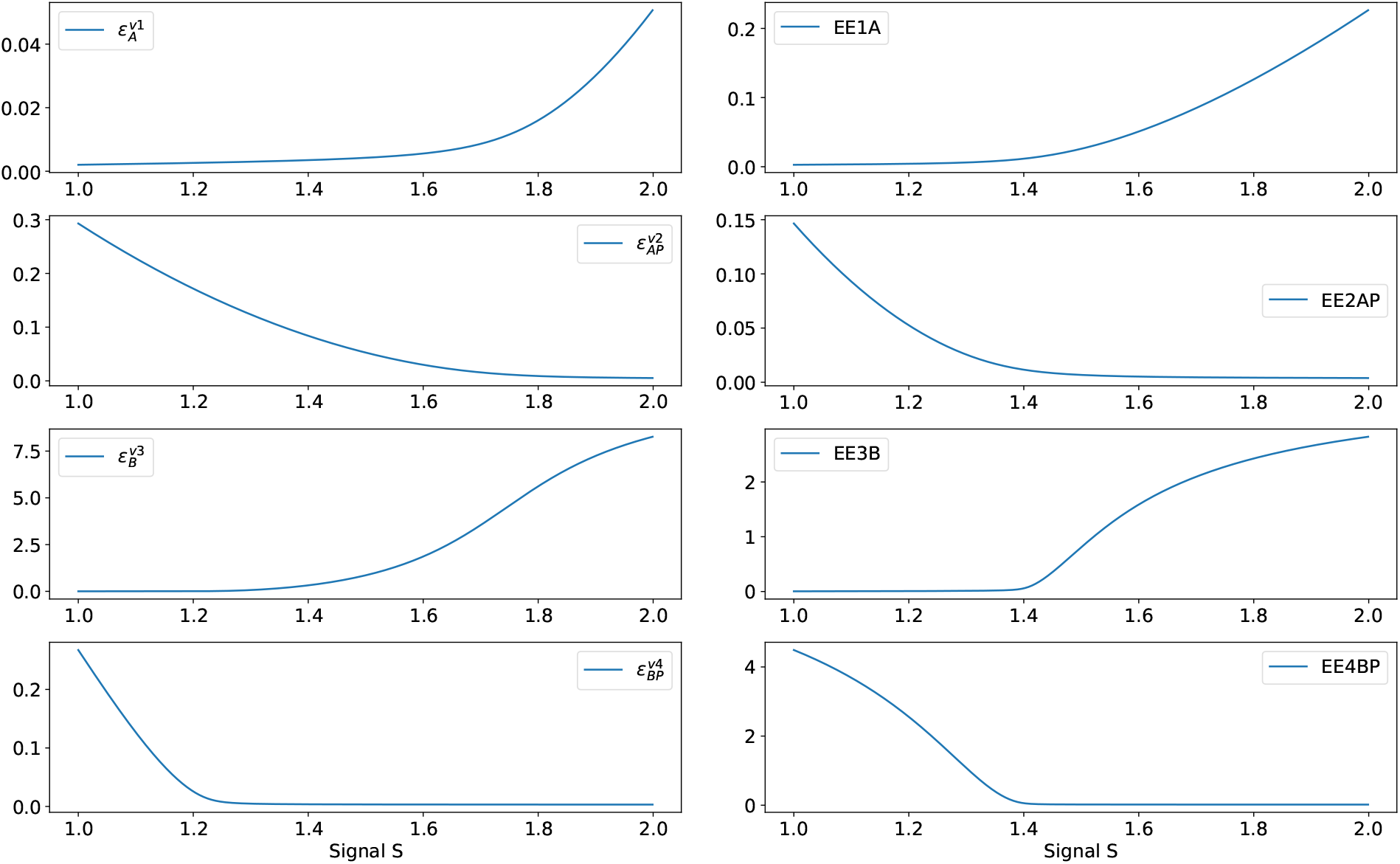
Elasticities involved the damping ratio for the two-layer one-cycle system. Left: unaligned, Right: aligned.

